# Efficient and specific oligo-based depletion of rRNA

**DOI:** 10.1101/589622

**Authors:** Amelie J. Kraus, Benedikt G. Brink, T. Nicolai Siegel

**Author notes:** To whom correspondence should be addressed: T. Nicolai Siegel, Biomedical Center Munich, Department of Physiological Chemistry, Ludwig-Maximilians-Universität München, Großhaderner Str. 9, 82152 Planegg-Martinsried, Germany.

## Abstract

In most organisms, ribosomal RNA (rRNA) contributes to >85% of total RNA. Thus, to obtain useful information from RNA-sequencing (RNA-seq) analyses at reasonable sequencing depth, typically, mature polyadenylated transcripts are enriched or rRNA molecules are depleted. Targeted depletion of rRNA or other highly abundant transcripts is particularly useful when studying transcripts lacking a poly(A) tail, such as some non-coding RNAs (ncRNAs), most bacterial RNAs and partially degraded or immature transcripts. While several commercially available kits allow effective rRNA depletion, their efficiency relies on a high degree of sequence homology between oligonucleotide probes and the target RNA. This restricts the use of such kits to a limited number of organisms with conserved rRNA sequences.

In this study we describe the use of biotinylated oligos and streptavidin-coated paramagnetic beads for the efficient and specific depletion of trypanosomal rRNA. Our approach reduces the levels of the most abundant rRNA transcripts to less than 5% with minimal off-target effects.

By adjusting the sequence of the oligonucleotide probes, our approach can be used to deplete rRNAs or other abundant transcripts independent of species. Thus, our protocol provides a useful alternative for rRNA removal where enrichment of polyadenylated transcripts is not an option and commercial kits for rRNA are not available.

## Introduction

Massive parallel sequencing has become the gold standard for transcriptome analyses and has been employed in organisms as diverse as humans ^1^, *Trypanosoma brucei* ^2^ and *Escherichia coli* ^3^. Yet, the large proportion of ribosomal RNA (rRNA), comprising >85% of total RNA in most organisms, complicates detection of low abundant transcripts. To increase the sequence coverage of the transcripts of interest, highly abundant transcripts, such as rRNAs, are typically removed from the pool of total RNA prior to sequencing ^4^.

Generally, one of two strategies is used to remove rRNA from total RNA, enrichment of mature polyadenylated (poly(A)) mRNA or targeted depletion of rRNA. The former is based on the use of oligo dT primers during reverse transcription of RNA into cDNA. This approach typically reduces the levels of rRNA to less than 5% ^5^, but it cannot be used if the RNA of interest is lacking a poly(A) tail. This is the case for partially degraded samples ^6^, many short and long ncRNAs ^7^, newly transcribed, unprocessed transcripts ^8^ or RNA from bacteria ^9^. To study these classes of RNA by RNA-seq, hybridization-based rRNA depletion is typically performed using one of the available kits that follow two approaches. One approach is to capture rRNA with complimentary oligos that are coupled to paramagnetic beads, followed by removal of the bound rRNA. Kits following this approach include for example the RiboMinus Transcriptome Isolation Kits (ThermoFisherScientific). An alternative strategy is to hybridize complementary DNA oligos to rRNA followed by degradation of the RNA:DNA hybrids using RNaseH ^10^, e.g. NEBNext rRNA Depletion Kits (New England Biolabs). To ensure efficient rRNA depletion and minimal removal of unrelated transcripts, both approaches require a high degree of sequence homology between rRNA transcripts and DNA probes. This restricts the use of the available commercial kits to organisms with rRNA sequences matching those of the provided probes. As a consequence, for many organisms no commercially available rRNA depletion kits exist. However, even if kits exist, the fact that oligo sequence and buffer composition are typically not disclosed poses problems to reproduce analyses if a manufacturer decides to change a kit’s composition or to terminate its production. For example, despite significant differences between human and *T. brucei* rRNA sequence, RiboMinus Eukaryote Kit (Invitrogen, A1083708) has been successfully used to deplete trypanosomal rRNA ^11,12^. However, following a change in the kit’s composition, the manufacturer is not recommending the kit for the depletion of trypanosomal rRNA (personal communication with ThermoFisherScientific). Similar problems arise following production stops. Even though a recent study comparing three kits for the depletion of bacterial rRNA found the Ribo-Zero kit (Illumina) to exhibit the highest efficiency ^13^, production of the kit has been discontinued as of November 2018.

Thus, our goal was to establish a protocol for the specific depletion of rRNA molecules that would allow the generation of high-quality transcriptome analyses without the need for commercially available kits.

Here we describe an efficient and highly specific rRNA removal approach that can be easily adapted to deplete rRNA or other transcripts of any species. Using a set of 12 biotinylated DNA oligos, we were able to reduce the levels of the most abundant rRNA transcripts to less than 5% of the total RNA with minimal off-target effects.

## Materials and Methods

### Cell cultivation and RNA isolation

Wild type *T. brucei* derived from the Lister 427 bloodstream-form MITat 1.2 isolate was cultivated at 37 °C and 5% CO2 in HMI-11 medium (HMI-9 medium ^14^ without serum plus) up to a density of 0.9 × 10^6^ cells/ml. 45 mio cells were harvested at 1,500 g and 4 °C for 10 min. The cell pellet was washed with 1 × TDB (5 mM KCl, 80 mM NaCl, 1 mM MgSO_4_, 20 mM Na_2_HPO_4_, 2 mM NaH_2_PO_4_, 20 mM glucose pH 7.4). RNA isolation was performed using the NucleoSpin RNA kit (Macherey-Nagel; cat. no. 740955.10) with minor changes. The cell lysis buffer was modified by adding 3.8 μl 1 M RNAse-free dithiothreitol (Sigma-Aldrich; cat. no. 10197777001) and 1 μl of 1:10 Ambion ERCC RNA Spike-In Mix (ThermoFisherScientific; cat. no. 4456739).

### Preparation of biotinylated oligonucleotides

Selected oligonucleotides were ordered with a 5′-biotin tag and HPLC-purified from Sigma-Aldrich. Each oligo was diluted to 100 μM in nuclease-free 10 mM Tris pH 8 (diluted from Ambion 1 M Tris pH 8; Invitrogen; cat. no. AM9856). The rRNA depletion mix was generated by combining equal volumes of each 100 μM oligonucleotide stock.

### Removal of ribosomal RNA

The set-up for the hybridization reaction is based on a published protocol ^15^. All solutions were kept free from nucleases. For each hybridization reaction, 2 μg of total RNA were mixed with 10 μl of formamide (SigmaAldrich; cat. no. F9037-100ML), 2.5 μl of 20 × SSC (3M NaCl, 0.3M sodium citrate, the pH was adjusted to 7.0 with HCl), 5 μl of 0.005 M EDTA pH 8 (stock solution 0.5 M; ThermoFisherScientific; cat. no. AM9260G), 2.48 μl of 100 μM rRNA depletion mix (total 4 μg of oligos) and RNAse-free water (ThermoFisherScientific; cat. no. AM9938) to a total volume of 50 μl. Hybridization was performed in a Bio-Rad C1000 Touch Thermal Cycler capable of performing temperature ramps with the following program: 5 min at 80 °C, ramp down to 25 °C at intervals of 5 °C per minute. Subsequently, 2 μl of RNAse-OUT (ThermoFisherScientific; cat. no. 10777019) and 50 μl of 1x SCC containing 20% formamide were added. Dynabead MyOne Streptavidin T1 beads (ThermoFisherScientific; cat. no. 65601) were prepared as recommended by the manufacturer for RNA applications and immobilization of nucleic acids. For each round of oligo capture 120μl (1200 μg) of magnetics beads were prepared (unless indicated otherwise). For the first round of depletion, the hybridization reaction was added to the beads, incubated at room temperature (RT) for 15 min using gentle rotation followed by bead separation on a magnetic rack (2 min). For the second and third round of depletion, the supernatant was added to a new batch of beads, incubated at RT for 15 min using gentle rotation followed by bead separation on a magnetic rack (2 min). The resulting supernatant, containing rRNA-depleted RNA, was purified using RNeasy MinElute CleanUp Kit (QIAGEN; cat. no. 74204). Depletion of rRNAs was evaluated on a 1.2% TBE-agarose gel and on an Agilent 2100 Bioanalyzer (Agilent Technologies; cat. no. G2939BA) using the RNA 6000 Nano Kit (Agilent Technologies; cat. no. 5067-1511).

### cDNA synthesis, library preparation and sequencing

Synthesis of cDNA was performed using NEBNext Ultra Directional RNA Library Prep Kit for Illumina (New England Biolabs; cat. no. E7420) according to manufacturer’s instruction. The concentration of cDNA was measured using Qubit dsDNA HS Assay Kit (Invitrogen, cat. no. Q32854) for Qubit 2.0 Fluorometer (Invitrogen; cat. no. Q32866). Sequencing libraries were prepared as described previously ^16^. To generate strand-specific RNA-seq libraries, uracil excision and removing of the second strand was performed prior to conversion of Y-shaped adapters. Therefore, 3 μl of USER enzyme (New England Biolabs; cat. no. M5505) were mixed with 16 μl of adapter-ligated DNA, 1 μl of TruSeq PCR primer cocktail (50 μM) and 20 μl of KAPA HiFi HotStart ReadyMix (KAPA Biosystems, cat. no. KK2601). USER digestion was performed at 37 °C for 15 min, followed by the published amplification protocol. Library concentrations were determined in duplicates using Qubit dsDNA HS Assay Kit (Invitrogen, cat. no. Q32854) for Qubit 2.0 Fluorometer (Invitrogen, cat. no. Q32866) and quantified using the KAPA Library Quantification Kit (KAPA Biosystems, cat. no. KK4824) according to manufacturer’s instruction. Strand-specific RNA-sequencing libraries were sequenced in paired-end mode on an Illumina NextSeq 500 sequencer 2 × 76 cycles.

### Processing of sequencing data

Adapter sequences were removed using Cutadapt ^17^ and the sequencing datasets were mapped to the *T. brucei* Lister 427 genome assembly (release 36, downloaded from TriTrypDB ^18^) using BWA-mem ^19^. The alignments were converted from SAM to BAM format, sorted and indexed using SAMtools version 1.8 ^20^. Additionally, unmapped, PCR or optical duplicate, not primary aligned and supplementary aligned reads were filtered out from the alignment files (SAM flag:3332).

The same procedure was applied for processing the poly(A) RNA-seq datasets from Hutchinson et al. ^21^.

### Differential expression analysis of RNA-seq data sets

Using BAM files, reads per gene were counted using the GenomicAlignments package ^22^ in R ^23^ and normalized to the total read counts. To determine the fraction of rRNAs in each RNA-seq experiment, rRNA read counts (28S alpha, 28S beta, 18S, 5.8S and 5S) were summed and their proportion of the total reads counts was calculated. Differential gene expression analyses were performed using the DESeq2 package ^24^ from R/Bioconductor. For analyses of non-rRNA transcripts, rRNA counts were excluded and vice versa.

### Comparison of ribo(-) and poly(A) RNA-seq analysis

The ribo(-) kit and different poly(A) RNA-seq datasets derived from previous studies were downloaded from NCBI SRA (accession number: SRP042959 ^12^) and EBI ENA (accession numbers: PRJEB8747 ^21^; PRJNA287144 ^25^; PRJEB22797^26^; PRJEB14403 ^27^). For correlation analyses, transcripts per million were calculated for each dataset in the non-alignment-based mode with SALMON ^28^ using the transcriptome of *T. brucei* Lister 427 (release 36, downloaded from TriTrypDB ^18^ as reference and normalized to the total number of transcripts excluding the number of rRNA transcripts.

### Calculation of reads mapping outside of ORFs

To calculate the number of reads mapping inside of protein-coding genes, we counted the number of reads mapping to the CDS features of the official annotation using the GenomicAlignments package ^22^ in R ^23^. Given the absence of UTRs in the annotation file of the *T. brucei* Lister 427 genome assembly (release 36, downloaded from TriTrypDB ^18^), we extended each CDS with the median UTR length of 89 bp upstream and 400 bp downstream ^2^. To calculate reads mapping outside of protein-coding genes, we counted all reads not mapping to CDS+UTRs or rRNA. The total number of reads mapping inside or outside of protein-coding genes was summed up and the percentages were calculated.

### Coverage visualization of RNA-seq data-sets

To visualize read coverage, filtered SAM alignments from replicate sequence experiments were merged using samtools *merge*. The number of reads was normalized per billion mapped reads and coverage files were generated in the wiggle format using COVERnant version 0.3.0 with the subcommand *ratio*, as previously described ^29^.

## Results

### Design of rRNA-specific biotinylated oligonucleotides

Just like ribosomes from other eukaryotes, the *T. brucei* ribosome contains more then 50 different rRNAs ^30^. Sequencing total RNA, we found that 28S alpha, 28S beta and 18S are the most abundant rRNA transcripts in *T. brucei*, contributing to 75% of the total *T. brucei* RNA (Supplementary Table 1). Thus, to deplete rRNAs of the small and large ribosomal subunits we established a protocol for the efficient removal of five rRNAs belonging to the major subunits (28S alpha, 28S beta, 18S, 5.8S and 5S) using customized, biotinylated hybridization probes and capturing of the probe:rRNA hybrid by streptavidin-coated beads (outlined in Fig. 1a). To ensure depletion of partially degraded transcripts, rRNA transcripts longer than 1 kb (28S alpha, 28S beta and 18S) were targeted with three probes against the 5′-end, the middle and the 3′-end of the transcript. The efficiency of rRNA removal strongly relies on sequence-specific hybridization of these probes and non-specific cross-hybridization can cause biases in transcriptome profiles.

To obtain highly specific probes, 25 bp probes were designed using Primer3Plus ^31^ and evaluated for possible cross-hybridizations using BLAST ^32^. Next, to increase the annealing temperature, the probe length was extended to 50 bp and re-evaluated for possible cross-hybridization using BLAST. A total of 12 probes (Table 1) were ordered with a 5′-biotin, to allow easy pull-out with streptavidin-coated paramagnetic beads.

**Table 1:**
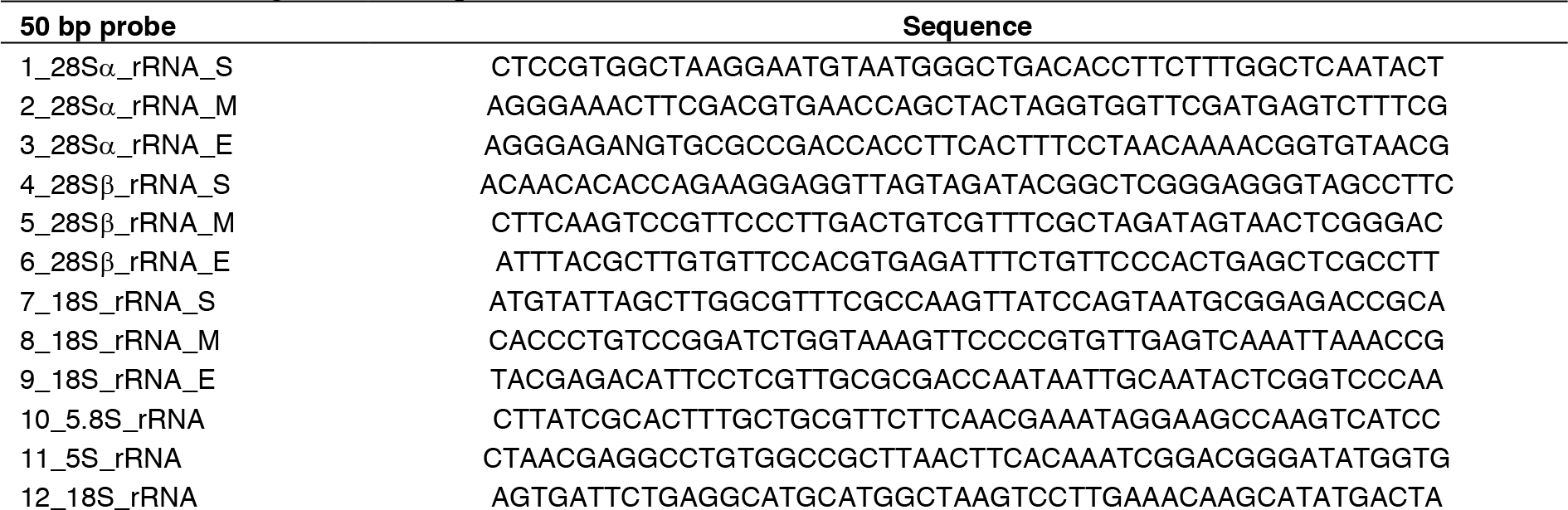
List of oligonucleotide probes.

### Using biotinylated oligos rRNA levels can be decreased to < 5% of total RNA

To establish conditions for efficient rDNA depletion, we compared different template-to-probe ratios and evaluated the effect of multiple rounds of rRNA depletion using different amounts of magnetic beads.

Previously, a template-to-probe mass ratio of 1:2 has been successfully used for the depletion of bacterial rRNA ^15^. Thus, we based our tests on similar ratios and compared hybridization of 2 μg of total *T. brucei* RNA with 0.5 μg, 1 μg, 2 μg or 4 μg (240 pmoles) of biotinylated oligos. Based on the binding capacity (provided by the manufacturer, ThermoFisherScientific) of the beads (Dynabeads MyOne Streptavidin C1) and the size of our oligos, 1.8 mg and 3.6 mg of beads seemed ideal to capture 2 μg and 4 μg of oligos, respectively.

Analyzing depleted and control RNA by gel electrophoresis, we observed the most efficient rRNA depletion when 4 μg of probes were used (Fig. 1b). In addition, our tests indicated that with 2 μg of total RNA and 4 μg of probes, three rounds of oligo capture using 1.2 mg of beads per round were necessary for the efficient depletion of rRNA (Fig. 1c).

Using these conditions, RNA-seq analysis of the rRNA-depleted and control samples indicated that following 0 (control), 1, 2 or 3 rounds of oligo capture the rRNA (28S, 5.8S, 18S and 5S) contributed to ~75%, ~30% and ~4% of total RNA, respectively (Fig. 1d and Supplementary Table 2). Thus, a set of 12 oligos was sufficient to reduce 28S alpha, 28S beta, 18S, 5.8S and 5S rRNA levels to < 5%.

**Figure 1.**
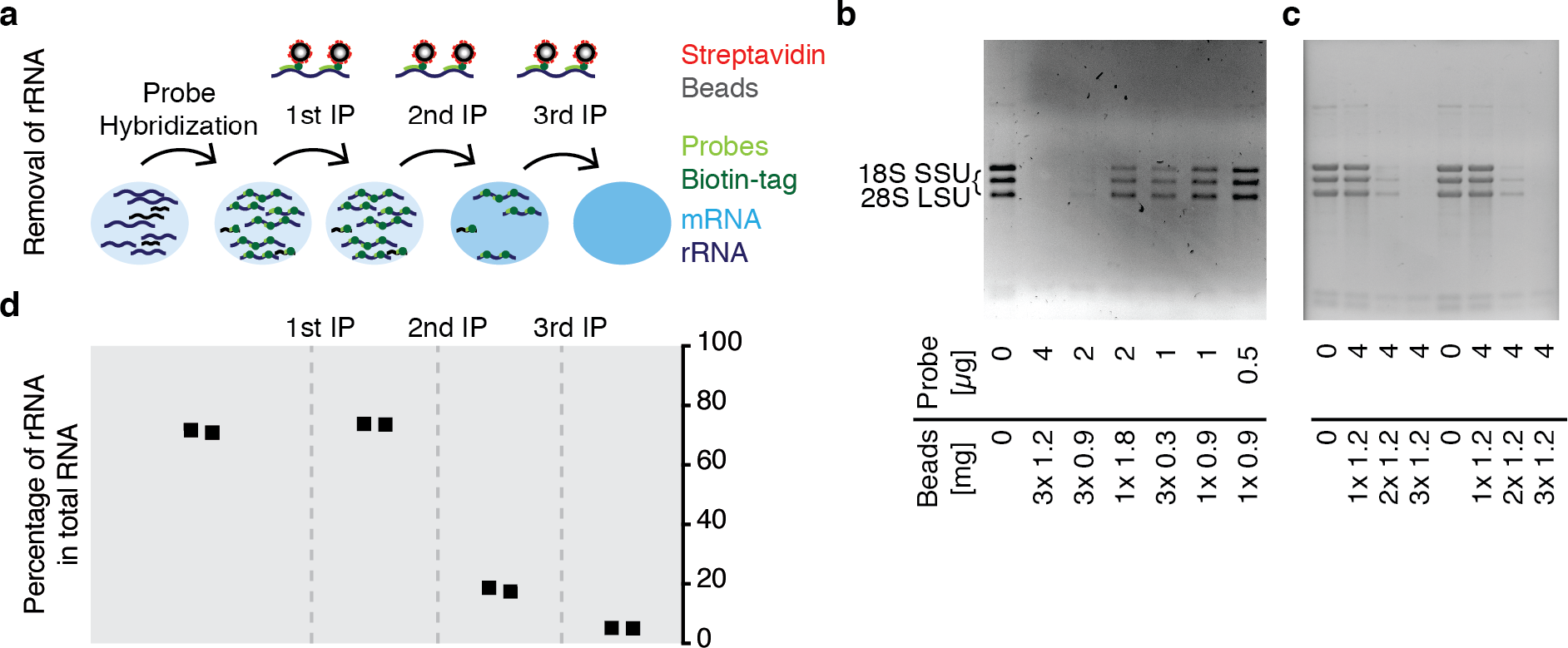
Depletion of rRNA transcripts. (a) Outline of depletion strategy. (b-c) Agarose gels revealing presence or absence of the three large trypanosomal rRNA transcripts following different depletion conditions. For each condition 2 μg of total RNA was used. (d) The samples shown in (c) were analyzed by RNA-seq. Shown are the percentages of 28S, 5.8S, 18S and 5S relative to total RNA.

### Depletion of non-rRNA is minimal

The usefulness of hybridization-based rRNA depletion strongly depends on its specificity as cross-hybridization of rRNA probes to other transcripts will result in incorrect transcript level measurements.

To determine whether the set of 12 biotinylated probes designed for this study cross-hybridizes with non-rRNA, we performed differential expression analyses, comparing the effect of different rounds of oligo capture with an un-depleted control sample. Analysis of untreated control RNA and RNA exposed to increasing rounds of oligo capture by RNA-seq, we found RNA levels of 0, 13 and 50 non-rRNA transcripts (of 8459 total) to be significantly (padj < 0.1) different by more than 2-fold after 1, 2 and 3 rounds of oligo capture (Fig. 2; supplementary Table 3). Twelve of the 13 transcripts affected by 2 rounds of oligo capture were also among the 50 transcripts affected by 3 rounds of oligo capture.

**Figure 2.**
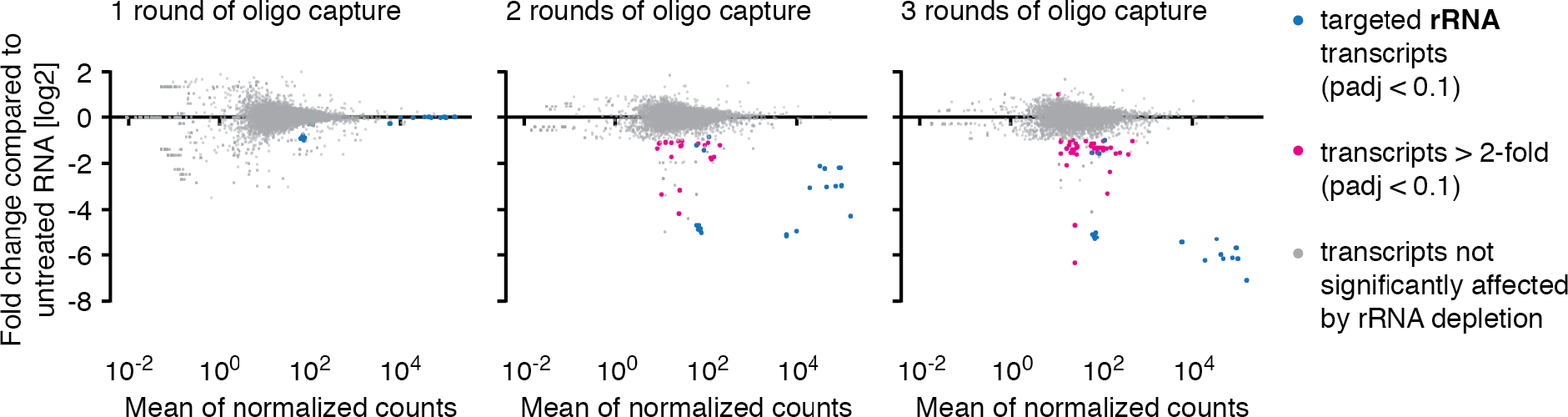
RNA-seq-based analysis of off-target depletion. Non-rRNA transcript levels were determined in duplicates by RNA-seq following 1, 2 or 3 rounds of oligo capture and compared to non-rRNA transcript levels of an untreated control sample. Reads mapping to rRNA genes were excluded from the mRNA off-target analysis and evaluated separately.

Given the reproducibility of off-target effects, we suspected the 50 co-depleted transcripts to share some homology with our rRNA-probes. Indeed, using BLAST ^32^, we observed for some of the co-depleted non-rRNA transcripts homology to our probes (Fig. 3 and Supplementary Table 4). Yet, other transcripts that exhibited a similar degree of homology showed no co-depletion. Overall, our data contains no evidence that the degree of homology between probe and non-rRNA transcript and the degree of co-depletion of non-rRNA transcripts are correlated.

Thus, while we were unable to determine why a set of 50 transcripts was co-depleted with rRNA transcripts, our rRNA-seq data indicated that 99.6% of genes are not affected by the depletion of rRNA transcripts.

**Figure 3.**
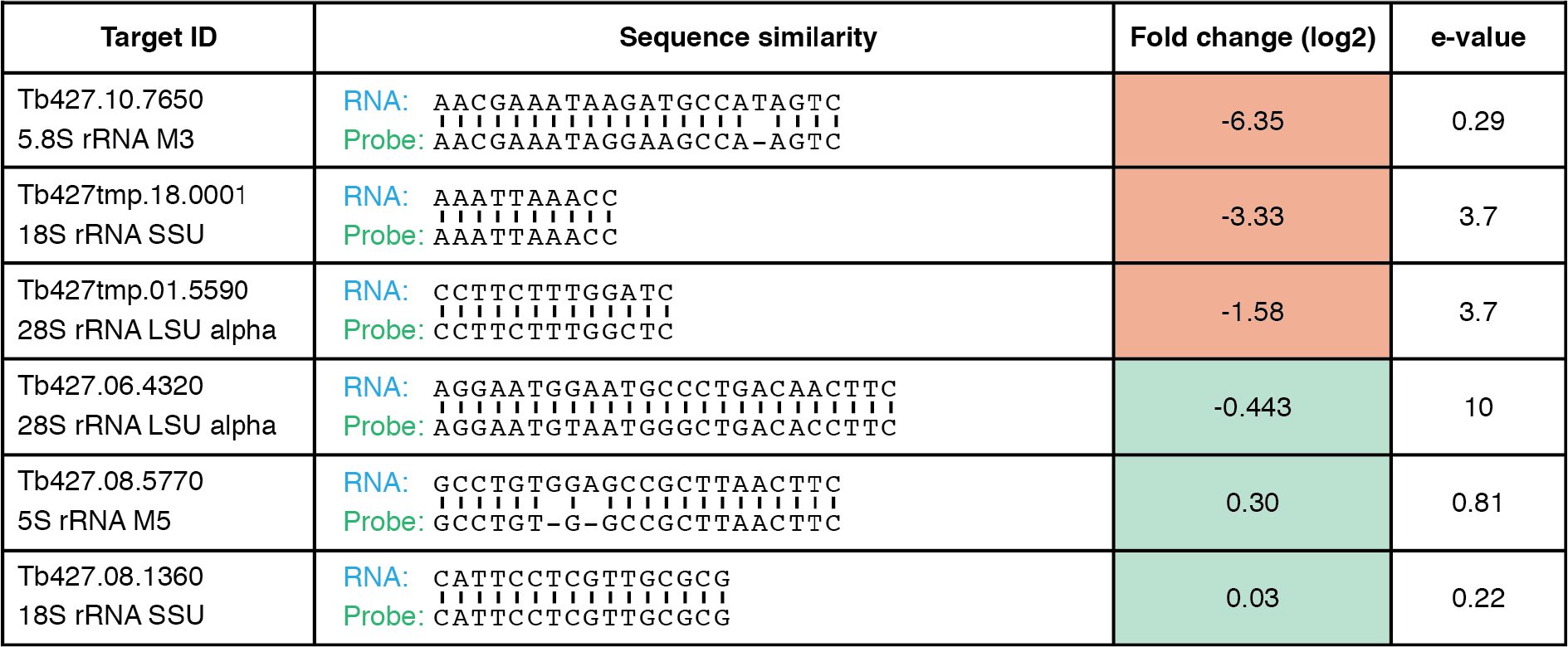
Representative examples of transcripts for which homology to an rRNA probe was observed and that were depleted (top three) or not depleted (bottom three) following rRNA depletion.

### rRNA depletion yields a higher percentage of ncRNA than poly(A) enrichment

Most transcriptome analyses benefit from the removal of rRNA. Yet the analysis of transcripts lacking a poly(A) tail, such as many non-protein coding transcripts and immature transcripts, eliminates the possibility of using strategies enriching for polyadenylated RNA. Instead, methods to specifically deplete rRNA are needed. Our RNA-seq data indicate that depletion of rRNA greatly improves the coverage across most genes (Fig. 4).

**Figure 4.**
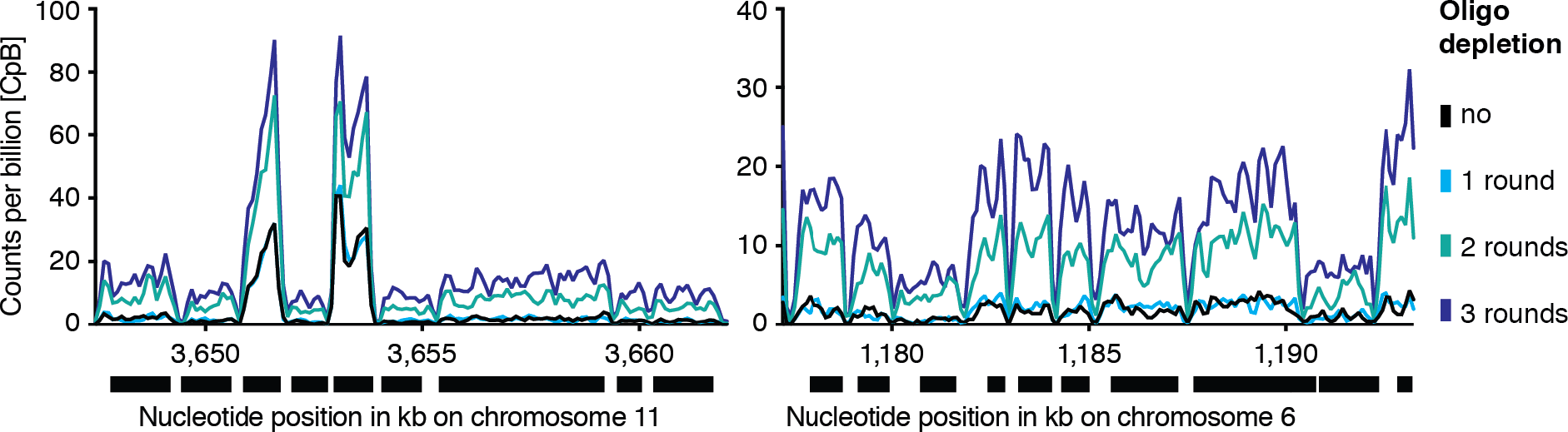
rRNA depletion affects RNA-seq coverage. Shown is the RNA-seq read distribution following multiple rounds of oligo capture across two representative genomic regions.

To better understand the effects of rRNA depletion and mRNA-enrichment-based approaches on transcriptome data, we compared the transcriptome data generated in this study with data from previously published mRNA-enriched RNA. We found the data reproducibility to be high with previously published RNA-seq datasets (Fig. 5 and Supplementary Table 5). Yet, for individual genes we observed marked differences in apparent transcript levels (Fig. 6). Overall, we found that transcriptome data from our rRNA-depleted pool of RNA contained a higher percentage of reads aligning outside of ORF than data from a poly(A)-enriched RNA pool: 79.8% compared to 54.5% ^21^. Such reads can be splicing intermediates, mRNA fragments mapping to the 5′or 3′UTR or ncRNA.

**Figure 5.**
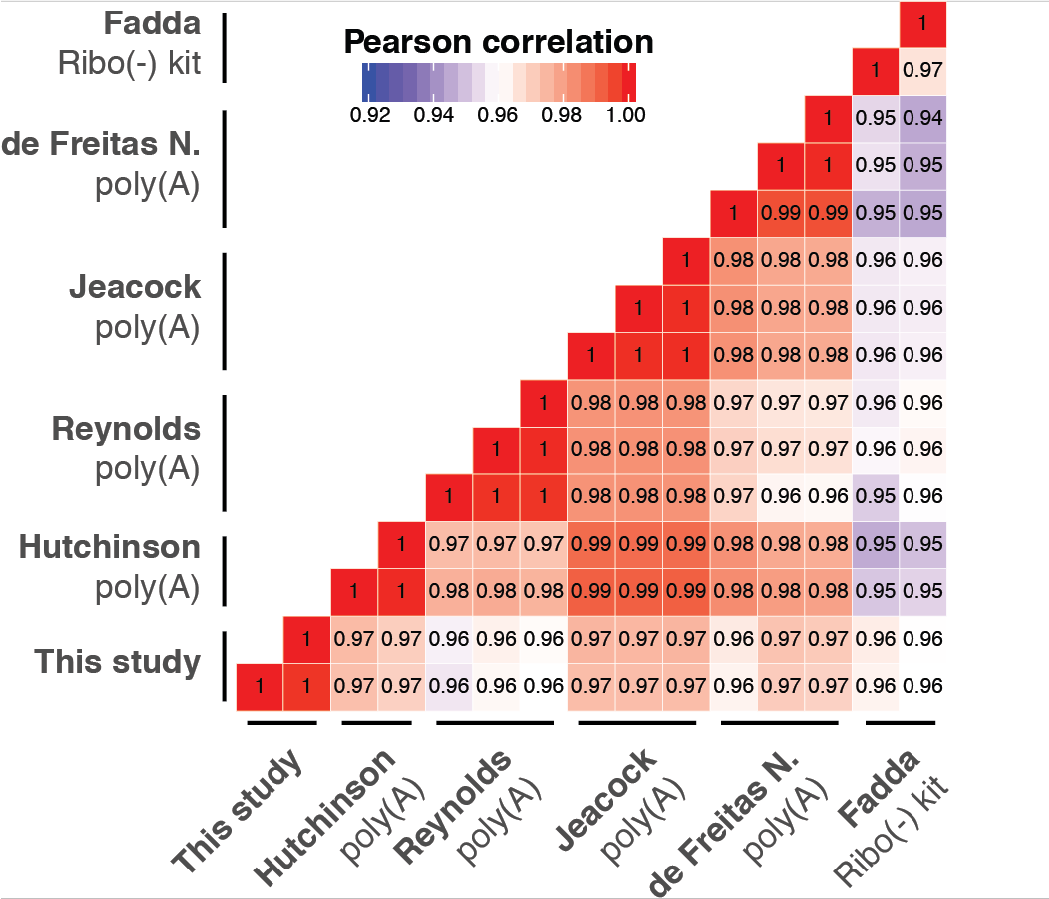
Correlation of different transcriptome analyses ^12,21,25–27^ and this study (3x oligo capture). Pearson coefficients were calculated based on the relative abundance of transcripts, excluding rRNA transcripts.

## Discussion

We describe the establishment of a highly specific and efficient approach to deplete rRNA from total RNA extracts that can be easily adopted to remove any species-specific rRNA or other abundant transcripts that may mask signal from low abundance transcripts.

Performing a systematic RNA-seq analysis following multiple rounds of rRNA depletion, we found that 3 rounds of probe capture were necessary to reduce rRNA levels to <5%. In addition, analysis of RNA-seq data revealed a small set of transcripts that was co-depleted with rRNA. Surprisingly, the degree of co-depletion did not correlate with the degree of sequence homology between the co-depleted transcripts and the probes. Thus, we suspect that the off-target effects may be caused by non-specific binding of some transcripts to the streptavidin-coated beads. Given that the off-target depletion was reproducible, it should be possible to normalize for the observed co-depletion if necessary. In addition, the reproducibility means that the co-depletion will not affect comparative RNA-seq analyses. If a specific transcript is co-depleted at similar levels in RNA from wild type and mutant cells, the co-depletion will not affect the measurements of changes in transcript level. Finally, we suspect that strategies employing enrichment of polyadenylated transcripts introduce biases as well. Differences in poly(A) tail length may very well affect the efficiency of poly(T)-primed reverse transcription ^33^.

To evaluate the robustness of our protocol, we compared the distribution of RNA-seq reads from our rRNA-depleted samples to previously published RNA-seq datasets derived from poly(A)-enriched samples. Comparing reads mapping to ORFs, we observed very high correlations, underscoring the reproducibility of the different methods. However, in our dataset a much higher percentage of reads mapped to regions outside of annotated ORF, to UTR or intergenic regions. We suspect this increase to be the result of the lack of enrichment for polyadenylated mRNA and to better represent the true distribution of cellular RNA. In this analysis we focused on the depletion of five rRNA transcripts. However, the *T. brucei* transcriptome contains several other highly abundant transcripts such as VSG-2 transcripts (4% of total RNA after 3 rounds of rRNA depletion) or rRNAs from the gamma, delta and zeta subunits (~10% of total RNA in our control sample). To increase the detection of low abundant mRNA even further, our set of 12 oligos may be extended to target other highly abundant transcripts.

**Figure 6.**
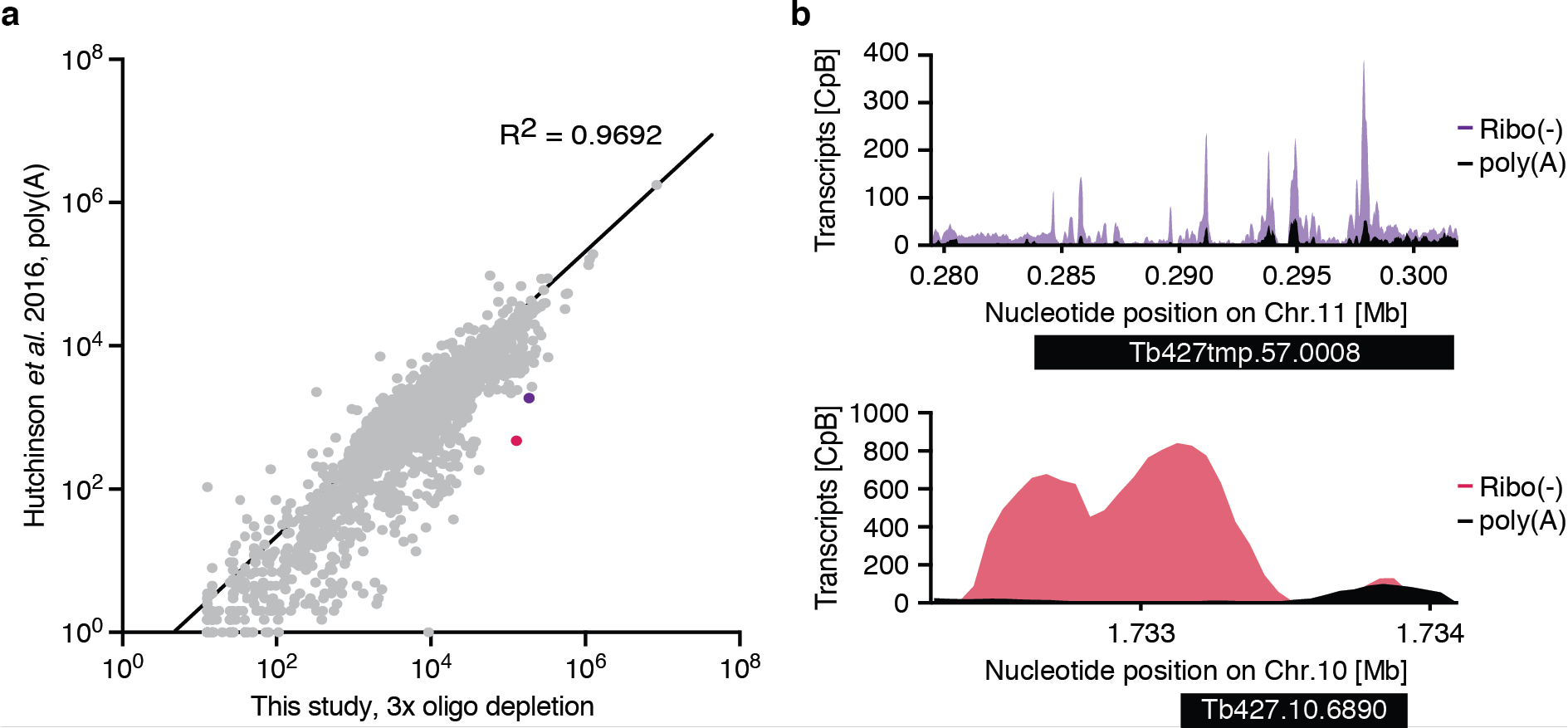
Comparisons of poly(A)-enriched and rRNA-depleted transcriptome datasets. (a) Scatter plot showing the normalized abundance of transcripts for data sets generated from poly(A)-enriched RNA 21 and our datasets generated from RNA following three rounds of oligo capture. Sequence tag abundance was normalized. (b) Examples of genes showing a different read coverage between a poly(A)-enriched dataset 21 and our dataset.

While the need for three rounds of depletion scientifically adds to the cost of our approach, the cost per rRNA depletion is still lower than it would be with most commercially available kits. To further reduce the costs, we propose regeneration of the streptavidin-coated magnetic beads. It has been shown that the incubation of beads in water at temperatures above 70 °C destroys the biotin-streptavidin interactions without denaturing the proteins ^34^. Alternatively, beads have been regenerated at 90% efficiency by exposure to a solution containing 25% aqueous ammonia and 25% aqueous ammonia in methanol (9:1) at 25 °C ^35^.

In summary, our study shows that organism-specific design of hybridization probes can be successfully used for the depletion of abundant transcripts. We describe a protocol that allows the efficient depletion of rRNA with little off-target effects. Thus, our protocol provides a useful alternative for the depletion of rRNA where enrichment of polyadenylated transcripts is not an option and commercial kits for rRNA are not available.

## Data availability

All sequencing data generated for this publication have been deposited in the European Nucleotide Archive and can be accessed through the accession number PRJEB31609. For details of the data analysis see Supplementary Table 6.

## Acknowledgements

We thank all members of the Siegel laboratory for valuable discussions. In addition, we thank Raúl Oscar Cosentino for his bioinformatic input and for critically reading the manuscript. We thank the Core Unit Systems Medicine of the University of Würzburg for the high-throughput sequencing. This work was funded by the Young Investigator Program of the Research Center for Infectious Diseases (ZINF) at the University of Würzburg, Germany, a grant from the German Research Foundation (SI 1610/2-1) and by an ERC Starting Grant (3D_Tryps 715466).

## Author Contributions

The study was conceptualized by AJ Kraus and TN Siegel. Experimental work was carried out by AJ Kraus, data analysis was performed by AJ Kraus and B Brink and the manuscript was written by AJ Kraus and TN Siegel.

